# Neural precursor cells rescue symptoms of Rett syndrome by activation of the Interferon γ pathway

**DOI:** 10.1101/2024.01.07.574507

**Authors:** Angelisa Frasca, Federica Miramondi, Erica Butti, Marzia Indrigo, Maria Balbontin Arenas, Francesca M. Postogna, Arianna Piffer, Francesco Bedogni, Lara Pizzamiglio, Clara Cambria, Ugo Borello, Flavia Antonucci, Gianvito Martino, Nicoletta Landsberger

## Abstract

The beneficial effects of Neural Precursor Cell (NPC) transplantation in several neurological disorders are well established and they are generally mediated by the secretion of immunomodulatory and neurotrophic molecules. We therefore investigated whether Rett syndrome (RTT), that represents the first cause of severe intellectual disability in girls, might benefit from an NPC-based therapy.

Using *in vitro* co-cultures, we demonstrate that, by sensing the pathological context, NPC-secreted factors induce the recovery of morphological and synaptic defects typical of *Mecp2* deficient neurons. *In vivo,* we prove that intracerebral transplantation of NPCs in RTT mice significantly ameliorates neurological functions. To uncover the molecular mechanisms underpinning the mediated benefic effects, we analysed the transcriptional profile of the cerebellum of transplanted animals, disclosing the possible involvement of the Interferon γ (IFNγ) pathway. Accordingly, we report the capacity of IFNγ to rescue synaptic defects, as well as motor and cognitive alterations in *Mecp2* deficient models, thereby suggesting this molecular pathway as a potential therapeutic target for RTT.

## Introduction

Mutations in the X-linked *MECP2* gene, encoding the methyl-CpG binding protein 2 (MeCP2), cause a broad spectrum of neuropsychiatric diseases including Rett syndrome (RTT), an early-onset neurodevelopmental disorder that represents the most common genetic cause of severe intellectual disability in girls worldwide (Amir et al., 1999; Chahrour and Zoghbi, 2007). RTT patients are diagnosed on the base of clinical signs, which include a period of regression together with the following four main criteria: loss of spoken language, loss of purposeful hand movements, gait abnormalities and the presence of hand stereotypies. Other clinical symptoms, such as cognitive disabilities, impairments in ambulation, breathing abnormalities and seizures, vary in frequency and severity, and emerge during the disease progression (Neul et al., 2010). Furthermore, deceleration in head growth, leading to microcephaly, typically appears within the first year of life (Armstrong, 2002).

Both cellular and animal models crucially contributed to expanding our knowledge on RTT and Mecp2 functions. Studies on neurons derived from either transgenic mice or human induced pluripotent stem cells (hiPSCs) highlighted morphological and functional alterations typically associated with *MECP2* mutations. Indeed, RTT neurons display reduced soma size and dendritic arborization, spine dysgenesis and decreased synapses’ number (Fukuda et al., 2005; Chapleau et al., 2009; Tropea et al., 2009; Belichenko et al., 2009; Jentarra et al., 2010; Ananiev et al., 2011; Perego et al., 2022). These defects contribute to altered calcium signaling and excitatory/inhibitory balance (Dani et al., 2005; Calfa et al., 2015; Bedogni et al., 2016). RTT symptoms are also well recapitulated by animal models mutated in *Mecp2*, among which the most used in RTT research is represented by the knock-out male mouse (*Mecp2* KO or null) (Guy et al., 2001; Chen et al., 2001). *Mecp2* null males appear grossly normal until 4-5 weeks of age, when they start manifesting progressive symptoms such as hind-limb clasping, ataxia, hypotonia, reduced movements, breathing and cognitive defects, seizures, and premature death (Guy et al., 2001; Chen et al., 2001; Cobolli Gigli et al., 2016). Although RTT mainly affects girls, male hemizygous animals remain largely used in basic research and pre-clinical studies, instead of female heterozygous mice. This is due to the fully penetrant phenotype of null male mice and the fact that female heterozygous animals manifest milder and delayed symptoms, with a higher variability, compared to the hemizygous model, because of random X-chromosome inactivation.

RTT brains are characterized by severe neurobiological changes, which are recapitulated by *Mecp2* null animals. Whereas neurodegeneration is absent, we recently reported that *Mecp2* deficiency causes delayed brain growth (Carli et al. 2023) and several studies described structural and functional abnormalities mainly affecting neurons, although with brain region specificity (Armstrong, 2005). These alterations are paralleled and caused by molecular abnormalities (Scaramuzza et al., 2021), and in line with the role of MeCP2 as transcriptional regulator, its loss of function causes subtle but widespread gene deregulation, as confirmed by several RNA sequencing analyses (Be-Shachar et al., 2009; Bedogni et al., 2016; Pacheco et al., 2017; Gogliotti et al., 2018; Sanfeliu et al., 2019). However, RTT does not spare glial cells that in fact are characterized by defective gene expression, morphology, and functionality (Ballas et al., 2009; Lioy et al., 2011; Yasui et al., 2013; Delépine et al., 2015; Pacheco et al., 2017; Albizzati et al., 2022). Consequently, Mecp2 defective astrocytes provide limited support to neuronal maturation and synapse formation (Albizzati et al., 2023; Sun et al., 2023). The scenario is further complicated by the presence of metabolic and mitochondrial abnormalities (Kyle et al., 2018; Shulyakova et al., 2017) and a dysregulation of immune response (Theoharides et al., 2014; Pecorelli et al., 2020), that likely play an active role in the generation and/or maintenance of RTT phenotypes. Evidence associating MeCP2 with the immune system is increasing, and cytokine dysregulation was described in RTT patients (Leoncini et al., 2015; De Felice et al., 2016; Byiers et al., 2020). Such a wide spectrum of symptoms challenges the identification of effective pharmacological therapies for RTT. Accordingly, despite several pre-clinical studies, success in clinical trials has been so far relatively poor (Frasca at al., 2023; Palmieri et al., 2023). However, very recently, Trofinetide, the synthetic version of the tripeptide glycine-proline-glutamate of IGF-1, was approved by FDA for RTT, considering its efficacy in improving core symptoms, including behaviour and communication (NCT04181723). Furthermore, two phase I/II clinical trials for gene therapy in RTT were recently authorized (Palmieri et al., 2023).

Considering all above, and the underlined difficulties in identifying valid therapeutic targets for the treatment of RTT, we tested the therapeutic potential of a cell-therapy, based on adult Neural Progenitor Cells (NPCs). NPCs constitute a class of multipotent stem cells, that has been widely studied for their ability to improve neuropathological signs in many neurodegenerative diseases characterized by spatially circumscribed CNS lesions, such as Parkinson’s disease and Huntington’s chorea, and also in multiple sclerosis, ischemia, and spinal cord injury (Lindvall and Kokaia, 2010). Importantly, safety of applying NPCs to patients affected by multiple sclerosis was recently approved (Genchi et al., 2023).

Despite initially predicted to act by replacing damaged cells, NPCs can also protect the pathological brain through multiple alternative mechanisms, which imply the interaction of NPCs with resident neural and immune cells. Transplanted NPCs can adapt their fate and functions to the receiving diseased CNS and modulate astrocytes, microglia, and inflammatory cells in response to pathological processes through paracrine mechanisms (bystander effects). Indeed, by releasing specific molecules (neurotrophic factors, reactive species, binding proteins, purines, or cytokines) and forming a close network that persists after administration, transplanted NPCs exert immunomodulatory or neuroprotective functions (Kokaia et al., 2012; Golgberg et al., 2015; Bacigaluppi et al., 2016; De Feo et al., 2017).

Other stem cells, such as mesenchymal cells, were widely used with therapeutic purposes in the field of neurodevelopmental and neurocognitive disorders, and a clinical trial on a small group of RTT patients treated with NPCs was reported on a Chinese journal (Liu et al., 2013; Nabetani et al., 2023). However, to the best of our knowledge, no study reports exhaustive results supporting the efficacy of NPCs in RTT. In this study, we tested our hypothesis that NPCs might exert positive effects on RTT models. *In vitro*, we prove that RTT neurons induce NPCs to secrete factors able to recover morphological and synaptic defects of null and heterozygous neurons. *In vivo*, we demonstrate that NPC transplantation ameliorates many RTT-like symptoms. Activation of the Interferon gamma–pathway (IFNγ) emerges as an involved mechanism, pointing to this cytokine as a novel and promising therapeutic target for RTT.

## Results

### By sensing the pathological context, NPCs secrete factors which promote neuronal maturation of *Mecp2* KO primary neurons

To determine whether NPCs exert beneficial effects on *Mecp2* deficient neurons, we initially used an *in vitro* system in which NPCs were seeded on transwell inserts and cultured with primary cortical neurons from DIV0 to the end of the experiment (DIV7 or DIV14) (**Fig 1A**). This co-culture allowed to assess the paracrine effects exerted by NPCs, in agreement with their well-known bystander mechanism (Kokaia et al., 2012). As controls we included WT and *Mecp2* KO neurons cultured with NIH3T3 fibroblasts, in addition to neurons cultured alone. Since RTT neurons suffer from a consistent reduction in dendritic arborization, we analysed dendritic complexity at DIV7, reporting the capacity of NPC-secreted factors to rescue both dendritic complexity and length in KO neurons (**Fig 1B-D**). In fact, while a significant reduction of the number of intersections between dendrites and concentric circles in Sholl analysis was measured both in KO neurons and in KO neurons cultured with fibroblasts (**Fig 1B,C**), a significant and general increment was observed in KO neurons treated with NPCs (**Fig 1C**). We reported a similar effect also in WT neurons (**Fig EV1A**), demonstrating the broad trophic effect of NPCs. Accordingly, by analysing the number of total intersections independently from distance to the soma, we found that the significant reduction in KO neurons was rescued when they were exposed to NPCs (**Fig EV1B**).

**Figure 1.**
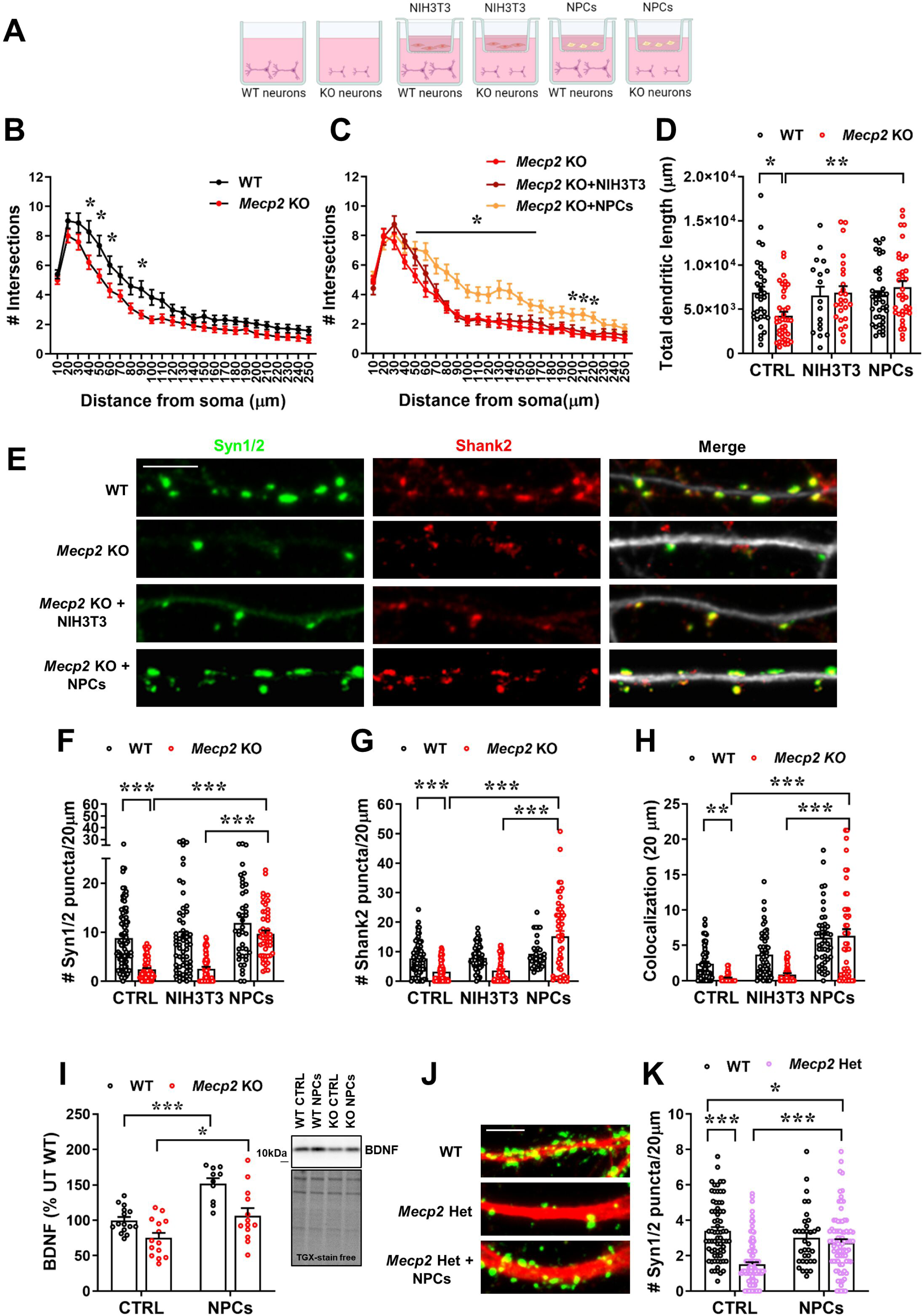
NPC treatment rescues dendritic branching and synaptic defects in *Mecp2* deficient neurons. **A)** The cartoon depicts the experimental setting, in which NPCs or NIH3T3 were seeded on transwell inserts, which were transferred on WT or *Mecp2* KO neurons at DIV0 until the end of the experiment (DIV7 for B-D; DIV14 for E-K). **B,C)** The graphs show the number of intersections calculated by Sholl analysis in WT and KO neurons (B), and in KO neurons cultured alone or with NIH3T3 or NPCs (C). **D)** The histogram indicates the total dendritic length calculated by NeuronJ in WT and KO neurons cultured alone (CTRL) or with NIH3T3 or NPCs. Data are reported as mean ± SEM. *p<0.05, **p<0.01 by two-way ANOVA, followed by Tukey post-hoc test. n=33 WT, n=17 WT+NIH3T3; n=39 WT+NPC; n=39 KO; n=26 KO+NIH3T3; n=32 KO+NPCs. Neurons derived from at least 3 different animals per genotype. **E)** Representative images of the immunostaining for Synapsin1/2 (Syn1/2; green), Shank2 (red) and Map2 (white), and the corresponding merge signal, of WT and KO neurons (DIV14) cultured alone or co-cultured with NIH3T3 or NPCs. Scale bar = 5 µm. **F-H)** Histograms indicate the mean ± SEM of number of Synapsin1/2 and Shank2 puncta in 20 µm (F,G) and of colocalized puncta (H) of WT and KO neurons cultured alone (CTRL) or co-cultured with NPCs or NIH3T3. **p<0.01, ***p<0.001 by two-way ANOVA, followed by Tukey post-hoc test; n=78 WT, n=58 WT+NIH3T3, n=41 WT+NPCs, n=68 KO, n=59 KO+NIH3T3, n=49 KO+NPCs. **I)** The histogram represents the mean ± SEM of the protein levels of the mature form of BDNF in WT and KO neurons cultured alone or co-cultured with NPCs (from DIV0 to DIV14), and expressed as percentage of WT neurons. *p<0.05, ***p<0.001 by two-way ANOVA followed by Tukey post hoc test. Representative bands of BDNF and the corresponding lanes of TGX-stain free gel are reported. **J)** Representative images of the immunostaining for Synapsin1/2 (Syn1/2; green) and Map2 (red) of WT and *Mecp2* Het neurons (DIV14) cultured alone or co-cultured with NPCs. Scale bar = 5 µm. **K)** The histogram indicates the mean ± SEM of Synapsin1/2 puncta density of WT and Het neurons cultured alone (CTRL) or co-cultured with NPCs. *p<0.05, ***p<0.001 by two-way ANOVA, followed by Tukey post-hoc test; n=68 WT, n=34 WT+NPCs, n=109 Het, n=83 Het+NPCs. Neurons derived from at least 6 different animals per genotype.

Importantly, NPC treatment also reverted the already described defects in synaptic puncta density and colocalization affecting KO neurons (Frasca et al., 2020). Indeed, by immunostaining for Synapsin1/2 and Shank2, we proved the ability of NPCs to rescue both alterations in KO neurons, while NIH3T3 cells revealed ineffective (**Fig 1E-H)**. In line with these effects and the established role of Brain Derived Neurotrophic Factor (BDNF) on synaptic plasticity, WB analysis indicated the already reported trend toward a reduction of Bdnf in KO neurons (Chang et al., 2016), that was recovered by NPC treatment (**Fig 1I**).

By analyzing *Mecp2* heterozygous (Het) neurons, that better recapitulate the human disorder, we confirmed the capacity of NPC-secreted molecules to increase the number of pre-synaptic puncta, without however completely rescuing the defect (**Fig J,K**). Interestingly, a significant reduction of the number of pre-synaptic puncta was found without observing a bimodal data distribution, indicating the already proved occurrence of non-cell autonomous mechanisms between neurons that express either the wild type or null *Mecp2* allele (Johnson et al., 2017; Belichenko et al., 2009). At the functional level, we analyzed spontaneous excitatory post-synaptic events (mEPSCs) by whole cell patch clamp recordings, revealing increased mEPSCs frequency in Het cultures with respect to WT neurons. This frequency alteration, that can be ascribed to several mechanisms, including an attempt of the system to compensate for the reduced pre-synaptic puncta, was fully rescued by NPCs but not by NIH3T3 cells. In addition, NPC secreted factors increased the mEPSC amplitude of Het neurons, which showed a slight and not significant reduction compared to WT cells (p=0.08) (**Fig EV1C**).

To explore the hypothesis that NPCs secrete factors depending on the specific environment (Drago et al., 2013), we analyzed the synaptic phenotype in KO neurons treated for 24 hours (DIV13-DIV14) with Conditioned Medium (CM) collected from the following co-cultures: WT neurons and NPCs (CM NPC^WT^), *Mecp2* KO neurons and NPCs (CM NPC^KO^), *Mecp2* KO neurons and NIH3T3 (CM NIH3T3^KO^). *Mecp2* KO and WT neurons cultured alone were used as controls (UT). Interestingly, the previously assessed synaptic defects that typically affect RTT neurons were exclusively rescued by the CM derived from NPCs co-cultured with KO neurons, indicating the capacity of the stem cells to sense the pathological environment and adapt their secretome in accordance with it, thereby releasing beneficial factors for RTT neurons (**Fig 2A-D**).

**Figure 2.**
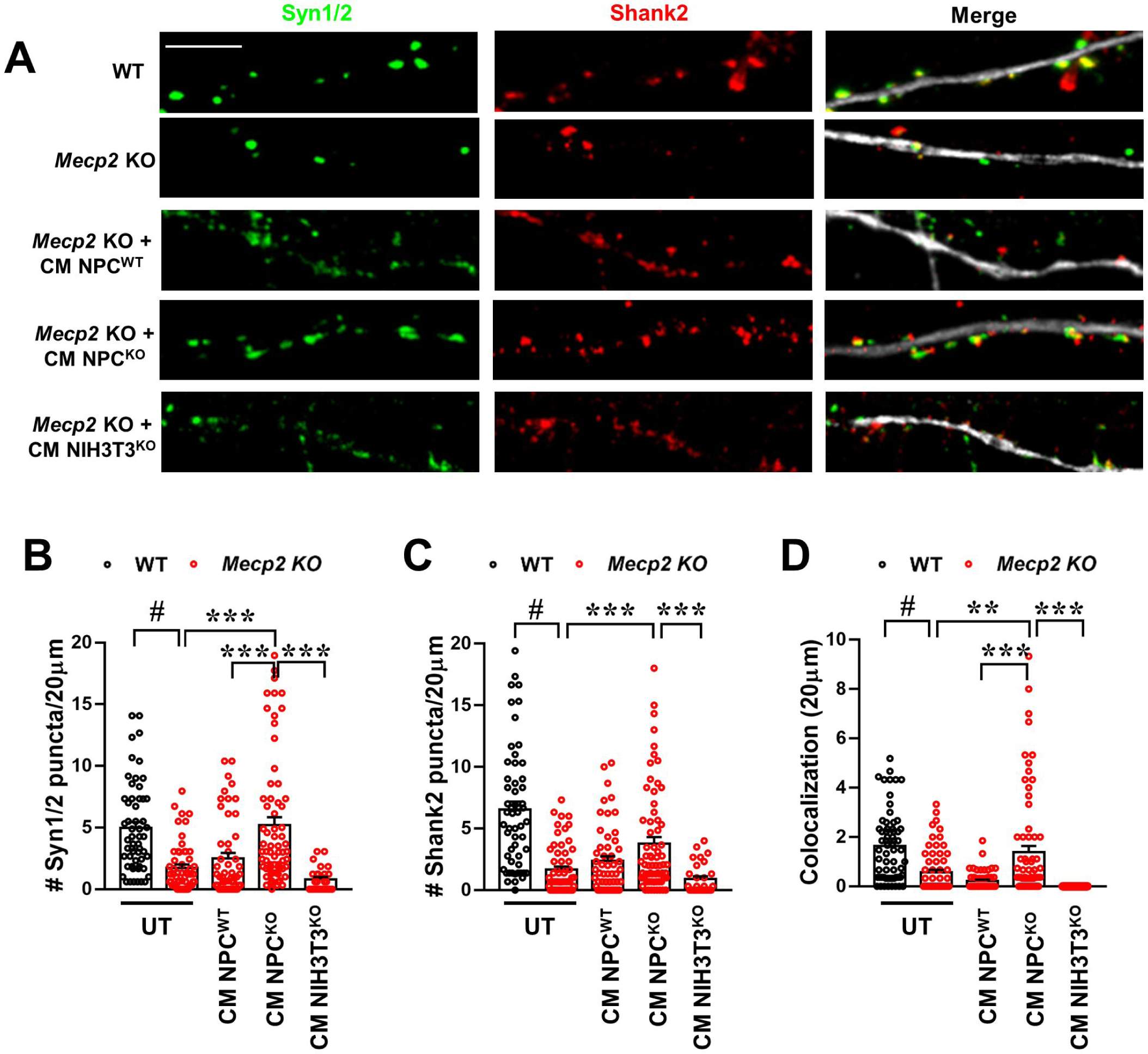
NPCs ameliorate synaptic defects in *Mecp2* KO neurons by sensing the pathological environment. **A)** Representative images of WT and *Mecp2* KO neurons (DIV14) immunostained for Synapsin1/2 (Syn1/2; green), Shank2 (red) and Map2 (white), left untreated (UT) or treated with the conditioned medium (CM) collected from the co-cultures between WT neurons and NPCs (CM NPC^WT^), or between *Mecp2* KO neurons and NPCs (CM NPC^KO^) or NIH3T3 (CM NIH3T3^KO^). Scale bar = 5 µm. **B-D)** Histograms show the mean ± SEM of Synapsin1/2 and Shank2 puncta density (B,C) and their colocalization (D). #p<0.001 by Mann Whitney test between WT and KO untreated neurons; **p<0.01,***p<0.001 by one-way ANOVA followed by Tukey post hoc test, among untreated *Mecp2* KO neurons and KO neurons treated with CM NPC^WT^, CM NPC^KO^ or CM NIH3T3^KO^. n=56 KO, n=51 KO+CM NPC^WT^, n=67 KO+CM NPC^KO^, n=28 KO+CM NIH3T3^KO^. WT and KO neurons derived from a pool of 3 different animals per genotype, whereas CM was collected from at least 6 different co-cultures per genotype.

### NPC transplantation ameliorates RTT-like symptoms in *Mecp2* deficient mice

To assess whether NPCs could improve RTT-related symptoms, 10×10^6^ cells were transplanted by intra-cisterna magna (i.c.m.) injection in symptomatic (P45-47) *Mecp2* KO mice, and WT littermates as control. Survival and behavioural parameters, which correlate with the severity of the disease (Guy et al., 2007; Cobolli Gigli et al., 2006; Scaramuzza et al., 2021), were analysed by a researcher blind to treatment and genotype, and data compared with PBS-injected WT/KO mice. Kaplan-Meyer survival analysis indicated a significant difference in the lifespan between NPC- and PBS-treated KO mice (**Fig 3B**). The well-defined scoring system was used to measure the severity of RTT-like symptoms in a period ranging between 5 days before and 16 days after transplant (Guy et al., 2007; Cobolli Gigli et al., 2006; Scaramuzza et al., 2021). Evolution of symptoms was graphically represented through a cumulative plot and a heatmap (**Fig 3C,D and Fig EV2A-E**), which shows that NPCs induced a widespread recovery of symptoms in KO animals starting roughly 10 days after transplantation. To investigate NPCs’ effectiveness in reverting neurological defects, animals were tested for their motor and cognitive functions. The rotarod test was used to assess both motor learning, comparing the latency to fall between the first trial with the subsequent trials, and motor coordination, analysing the latency to fall in the last trial (Buitrago et al., 2004). As expected, PBS-treated KO mice manifested impaired motor learning, which was significantly ameliorated by NPCs (**Fig 3E**), and analysis of motor coordination proved that the significant difference between WT and KO mice was completely rescued by NPC administration (**Fig 3F**). Furthermore, by performing the Novel Object Recognition (NOR) test, that evaluates the difference in the exploration time of a novel and a familiar object (expressed as Discrimination Index; D.I.), we reported that the significant defect in KO animals was no more present following NPC transplantation (**Fig 3G**). The amelioration observed in the NOR test was purely cognitive and not influenced by mobility, since the impairment in the distance travelled inside the arena was not affected by NPC treatment (**Fig 3H, EV2F**).

**Figure 3.**
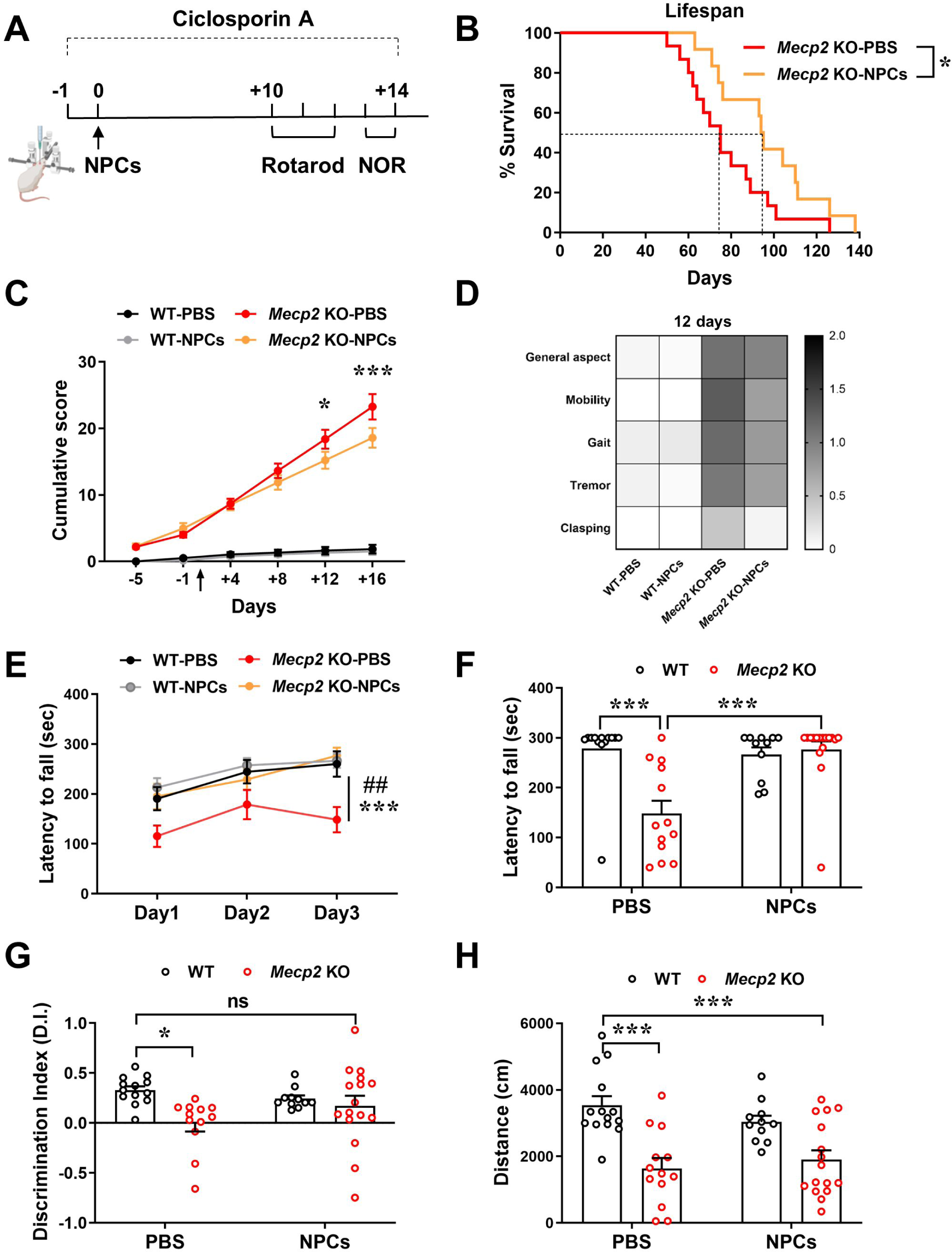
NPC transplantation prolongs the lifespan and ameliorates RTT-like impairments in *Mecp2* KO mice. **A)** NPCs were injected in WT and *Mecp2* KO mice (P45-47) and behavioural tests were conducted starting from 10 days after transplantation. Ciclosporin A (50 mg/Kg, s.c.) was daily administered both in PBS- and NPC-treated WT/KO animals starting the day before surgery and for 15 days. **B)** Kaplan-Meyer survival analysis shows that NPC transplantation prolongs the lifespan of KO mice compared to PBS-treated KO animals. The median survival corresponds to 70 days for PBS-KO mice and 94 days for NPC-KO animals. *p<0.05 by Gehan-Breslow-Wilcoxon test. n=13 PBS-treated KO, n=11 NPC-treated KO. **C)** The graph depicts the mean ± SEM of the cumulative phenotypic score calculated for all the experimental groups. The black arrow indicates the day of transplantation. Asterisk denotes a significant difference between NPC-KO and PBS-KO animals. The difference between PBS-KO and PBS-WT mice, or NPC-KO and PBS-WT is omitted, although significant at all time-points, excluding at day −5. *p<0.05, ***p<0.001 by two-way ANOVA followed by Tukey post hoc test. **D)** Heatmap of the phenotypic score at the 12^th^ day after NPC transplantation, indicating in a grey scale the severity for each symptom (from 0 = absent to 2 = very severe). n=15 WT-PBS, n=15 WT-NPCs, n=8 KO-PBS, n=12 KO-NPCs. **E)** The graph represents the mean ± SEM of the time (in seconds) spent on the rod during each day of the test, thus deducing motor learning. Asterisk denotes a significant difference between NPC-KO and PBS-KO animals, whereas hashtag indicates a significant difference between PBS-KO and PBS-WT mice. ## p<0.01, *** p<0.001 by two-way ANOVA followed by Tukey post hoc test. **F)** The histogram shows the mean ± SEM of the time (in seconds) spent on the rod at the 3^rd^ day of the test. ***p<0.001 by two-way ANOVA followed by Tukey post hoc test. **G)** The histogram represents the mean ± SEM of the discrimination (D.I.) index values, defined as follow: (time exploring the novel object – time exploring the familiar object) / total time) assessed by novel object recognition (NOR) test. *p<0.05 by two-way ANOVA followed by Tukey post hoc test; ns indicated no statistically significant difference between NPC-treated KO and PBS-treated WT animals. **H)** The histogram shows the distance (in cm) travelled during the first day of NOR test, expressed as mean ± SEM. ***p<0.001 by two-way ANOVA followed by Tukey post hoc test. In E-H, n=14 WT-PBS, n=12 WT-NPCs, n=11 KO-PBS, n=16 KO-NPCs.

To start characterizing the molecular mechanisms by which transplanted NPCs exert positive effects in KO mice, we investigated NPC grafting, localization, and differentiation. GFP staining revealed that most transplanted cells distributed posteriorly, in an area proximal to subarachnoid space of cerebellum, fourth ventricle and brainstem, in accordance with site of injection (**Fig 4A,B**). 10 days after transplantation we estimated that in the KO brain persisted an average of 200,000 GFP^+^-NPCs, corresponding to ∼ 2% of transplanted cells. Grafted NPCs expressed the stem cell marker Nestin, and most of them were positive for the astrocytic marker GFAP. However, the lack of a ramified astrocytic morphology suggested that these cells maintain an immature phenotype, probably committed to the astrocytic lineage (**Fig 4C**). These results led us to hypothesize that, similarly to the *in vitro* observations, transplanted NPCs could induce a beneficial action by sensing pathological context and releasing soluble factors, therefore through a bystander mechanism.

**Figure 4.**
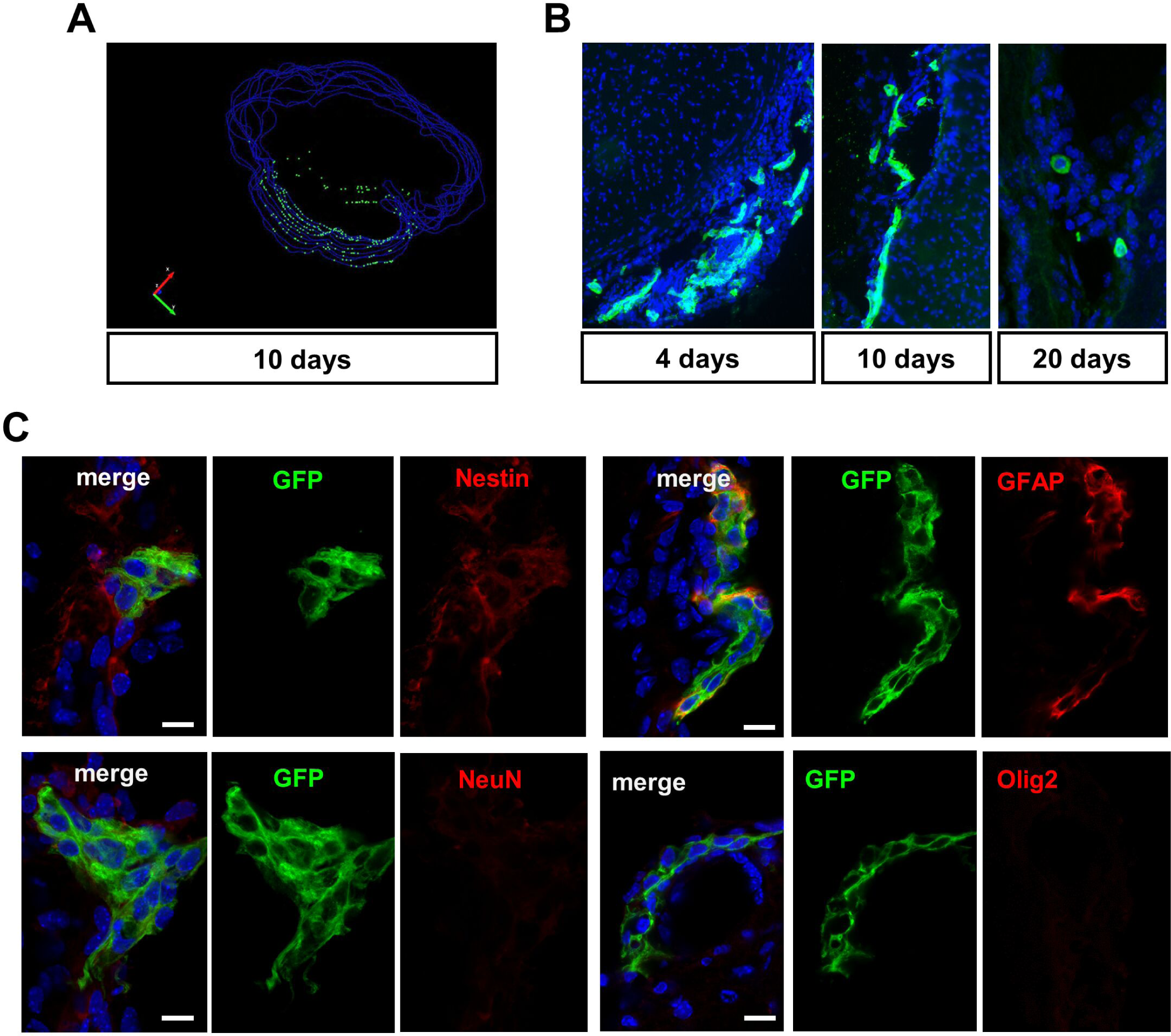
NPCs localize along the meninges in *Mecp2* KO brain and mainly retain an undifferentiated phenotype. **A)** 3D reconstruction of NPCs’ distribution in the KO brain by Neurolucida software**. B)** Representative images of GFP^+^-NPCs (green), localized along the meninges in the caudal region of the brain at 4, 10 and 20 days after transplantation. Nuclei are immunostained with DAPI (blue). **C)** Representative images of GFP^+^-NPCs (green) with different cell markers (red). Scale bar = 20 µm.

NPC grafting was also evaluated in P180 *Mecp2* Het mice, when symptoms are comparable to those observed in P45 KO males (Carli et al., 2023). Immunofluorescence analysis of GFP^+^ NPCs, performed 10 days after transplantation, revealed the presence of only few transplanted cells caudally localized, as observed in KO brains (**Fig 5A**). In possible good accordance with the limited grafting, assessment of phenotypic score and motor functions indicated that NPC treatment was not effective. Despite that, NOR test revealed the ability of NPCs to abrogate the memory defect that distinguishes untreated Het mice from WT, therefore supporting the therapeutic effects of these cells (**Fig 5B-D**).

**Figure 5.**
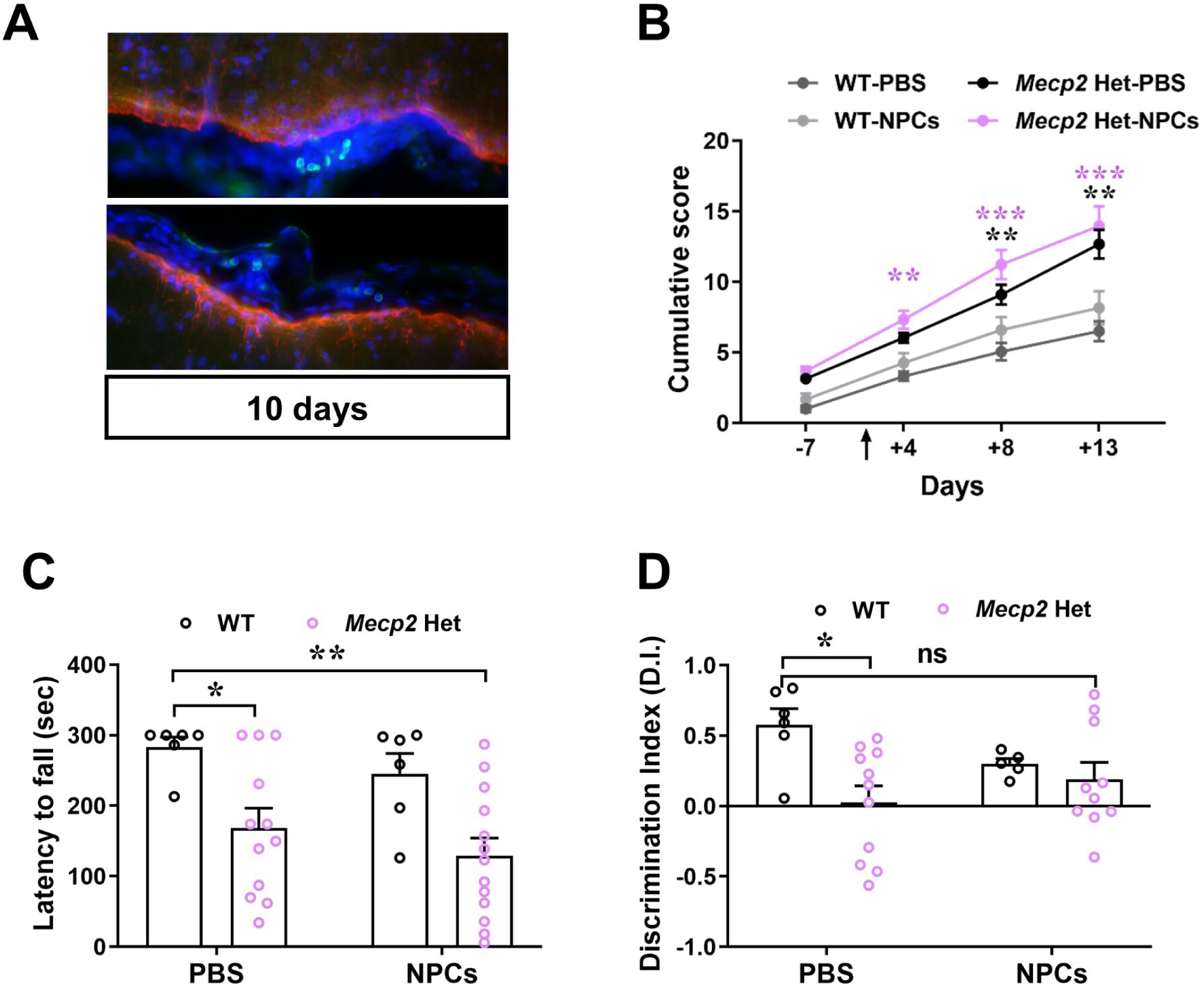
NPC transplantation in *Mecp2* Het animals only improve their cognitive defects, without ameliorating their general well-being and motor abnormalities, in accordance with their limited grafting. **A)** Representative image of GFP^+^-NPCs (in green) and GFAP (in red) in the caudal part of the Het brain 10 day after transplantation. Very few cells are present in the meninges of brainstem. **B)** The graph depicts the mean ± SEM of the cumulative phenotypic score calculated for female WT and *Mecp2* Het mice, injected with NPCs at P180. The black arrow indicates the day of transplantation. Black asterisk denotes a significant difference between PBS-Het and PBS-WT animals, whereas violet asterisk indicates a significant difference between NPC-Het and PBS-WT mice. **p<0.01, ***p<0.001 by two-way ANOVA followed by Tukey post hoc test. n=8 WT-PBS, n=6 WT-NPCs, n=11 Het-PBS, n=11 Het-NPCs. **C)** The graph represents the mean ± SEM of the time (in seconds) spent on the rod during the 3^rd^ day of the test. *p<0.05, **p<0.01 by two-way ANOVA followed by Tukey post hoc test. **C)** The histogram represents the mean ± SEM of the discrimination (D.I.) index values, assessed by novel object recognition (NOR) test. *p<0.05 by two-way ANOVA followed by Tukey post hoc test; ns indicated no significant difference between NPC-treated Het and PBS-treated WT animals. n=6 WT-PBS; n=5 WT-NPCs; n=12 Het-PBS; n=13 Het-NPCs.

### NPCs induce the deregulation of immune-related molecular pathways and the activation of IFNγ signalling

Bulk RNA-seq of WT and *Mecp2* KO cerebellum was performed to reveal the molecular mechanisms driven by the treatment. This cerebral region was selected considering the NPC distribution after transplantation and for its roles in motor and cognitive functions (Koziol, 2014) that were improved by NPCs. PCA analysis shows a clear separation between WT and *Mecp2* KO cerebella, while NPC transplanted samples are more similar to their controls (**Fig 6A).** Using a fold change >1 or <-1 as cut off for differential expression, we identified only a small percentage of DEGs (∼23%) differentiating KO from WT samples, therefore confirming that the lack of Mecp2 subtly modifies gene transcription (Sanfeliu et al., 2019) (**Fig 6B**). Using the same filters and comparing DEGs in transplanted KO tissues with respect to the untreated ones, we revealed a high number of DEGs; interestingly, genes with the higher entity of deregulation were upregulated (55 DEGs with a fold change >1). No transcriptional change was observed in NPC-injected WT samples, in good accordance with the lack of behavioural effects of NPCs on WT mice (**Fig 6B**).

**Figure 6.**
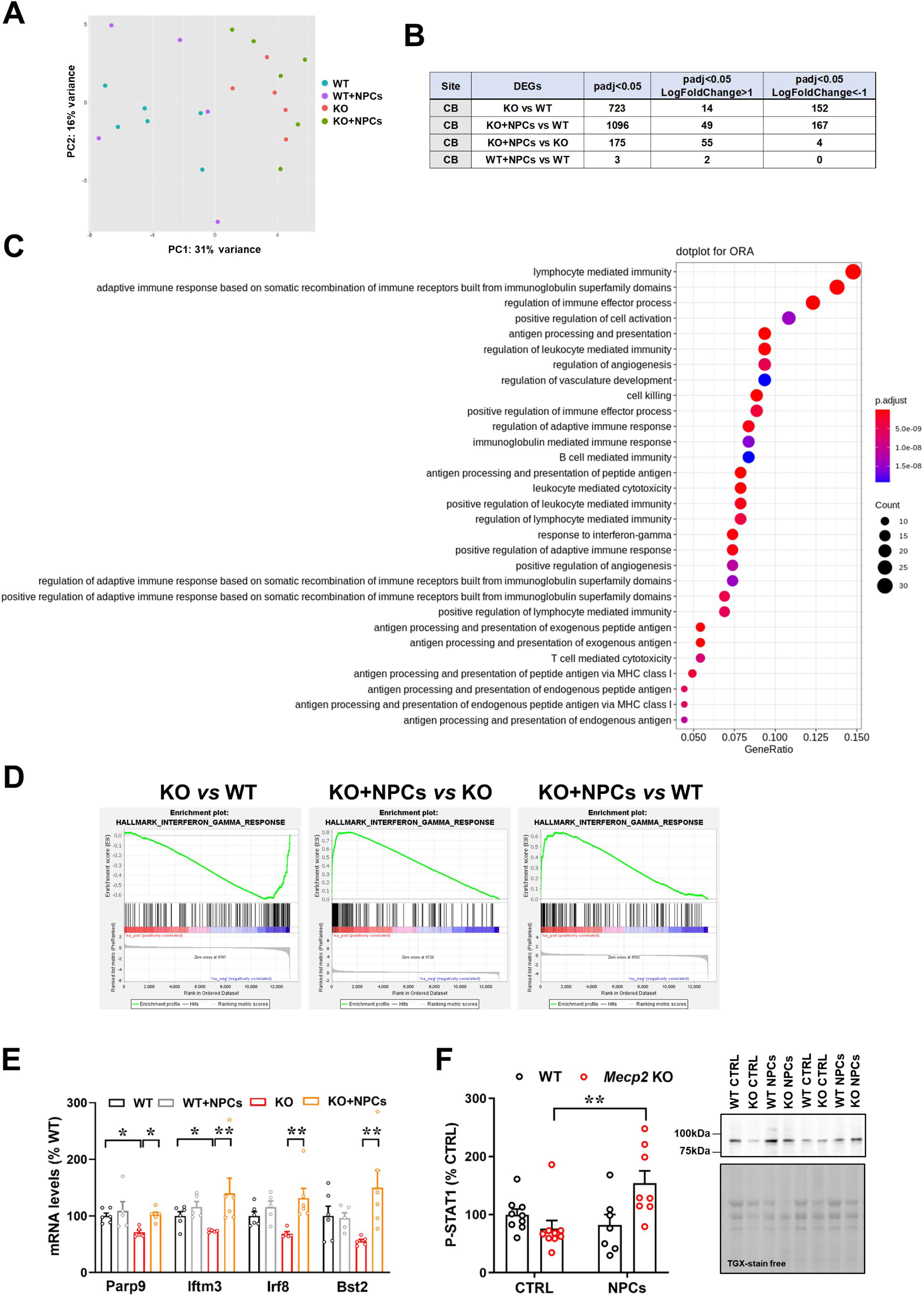
RNA sequencing revealed the activation of IFNγ pathway in the transplanted *Mecp2* KO cerebellum. **A)** Principal component analysis (PCA) plot for the sequenced samples of WT, *Mecp2* KO, WT+NPCs and *Mecp2* KO+NPCs cerebella. Percentage of variance is reported for both PC1 (first component) and PC2 (second component). **B)** The number of deregulated genes (DEGs) for the different comparison is reported, considering a p.adj<0.05. The number of DEGs with a LogFoldChange lower than −1 (down-regulated genes) or LogFoldChange greater than 1 (up-regulated genes) is also indicated. **C)** Dot plot of Gene Ontology (GO) enriched pathway analysis in the cerebellum, indicating the top 30 most enriched pathways of the comparison between KO+NPCs *versus* KO samples. **D)** Gene set enrichment analysis (GSEA) of IFNγ response in cerebellar samples, indicating a significant enrichment of the gene set in KO+NPCs vs KO comparison, as well as in KO+NPCs vs WT comparison. **E)** The histogram reports the transcriptional levels of genes (Parp9, Iftm3, Irf8 and Bst2) associated to IFNγ pathway. Data, expressed as percentage of WT animals, are shown as mean ± SEM. *p<0.05, **p<0.01 by two-way ANOVA followed by Tukey post hoc test. **F)** Western blot analysis of phosphorylated STAT1 (P-STAT1) in WT or KO neurons cultured with NPCs (for 14 days) (n=7-9). Data are represented as mean ± SEM and expressed as percentage of WT neurons cultured alone. Representative bands of P-STAT1 and the corresponding lanes of TGX-stain free gel are reported.

To identify the biological pathways enriched in the list of DEGs resulting from the different comparisons, we performed Over Representation Analysis (ORA) (Boyle et al., 2004). The ORA results of the comparisons *Mecp2* KO-PBS *versus* WT-PBS revealed the deregulation of several pathways related to RTT dysfunctions, such as synapse organisation, ion channel transport and regulation of neurogenesis, thereby validating the RNA-seq analysis (**Fig EV3**) (Ben Shachar et al., 2009; Gogliotti et al., 2018; Sanfeliu et al., 2019).

Interestingly, the results shown in **Fig 6C**, in which we analysed the effects of NPCs on KO cerebella, revealed the deregulation of several pathways associated with immune response, including the pathway related to interferon gamma (IFNγ). Moreover, we took advantage of another approach to analyse gene expression profiles, i.e. gene set enrichment analysis (GSEA) (Subramanian et al., 2005) and we found that IFNγ response was less represented in the KO cerebellum compared to WT (ES=-0.64; q-value=0.001), whereas an enrichment was evident in KO-NPCs *versus* untreated KO (ES=0.80; q-value<0.001) and WT (ES=0.64; q-value<0.001), thus confirming ORA data (**Fig 6D**). By qRT-PCR we analysed the expression of a selected panel of genes associated with the IFNγ response, including Ppar9, Ifitm3, Irf8 and Bst2, validating the activation of this molecular pathway (**Fig 6E**). Coherently, NPCs co-cultured with KO neurons (but not with WT) secreted factors which increased the levels of phosphorylated Stat1, that is considered a good marker of activation of IFNγ pathway (**Fig 6F**).

### The cytokine IFNγ holds therapeutic potential for the treatment of RTT

Instructed by these molecular results, which pointed to the ability of NPCs to activate the IFNγ pathway in cellular and animal models of RTT, and encouraged by previous evidence indicating the neuroprotective effects mediated by the cytokine (Filiano et al., 2016), we tested potential benefits of an IFNγ administration. We thus exposed cultured RTT neurons to IFNγ and investigated its ability to rescue synaptic defects. Three doses (25, 75 and 100 ng/ml) were initially supplemented to medium, revealing that none of these was toxic either for WT or KO neurons (**Fig EV4A**). WB analysis confirmed that all the concentrations were able to induce Stat1 phosphorylation (**Fig EV4B**). Notably, by immunofluorescence, we proved that the cytokine reverted both the density of synaptic markers and their colocalization in KO neurons in a dose-dependent manner (**Fig 7A-D**). The highest dose of IFNγ reverted also the defective density of pre-synaptic puncta in Het neurons (**Fig 7E, F**). No detrimental effect was observed in WT neurons (**Fig 7B-F**).

**Figure 7.**
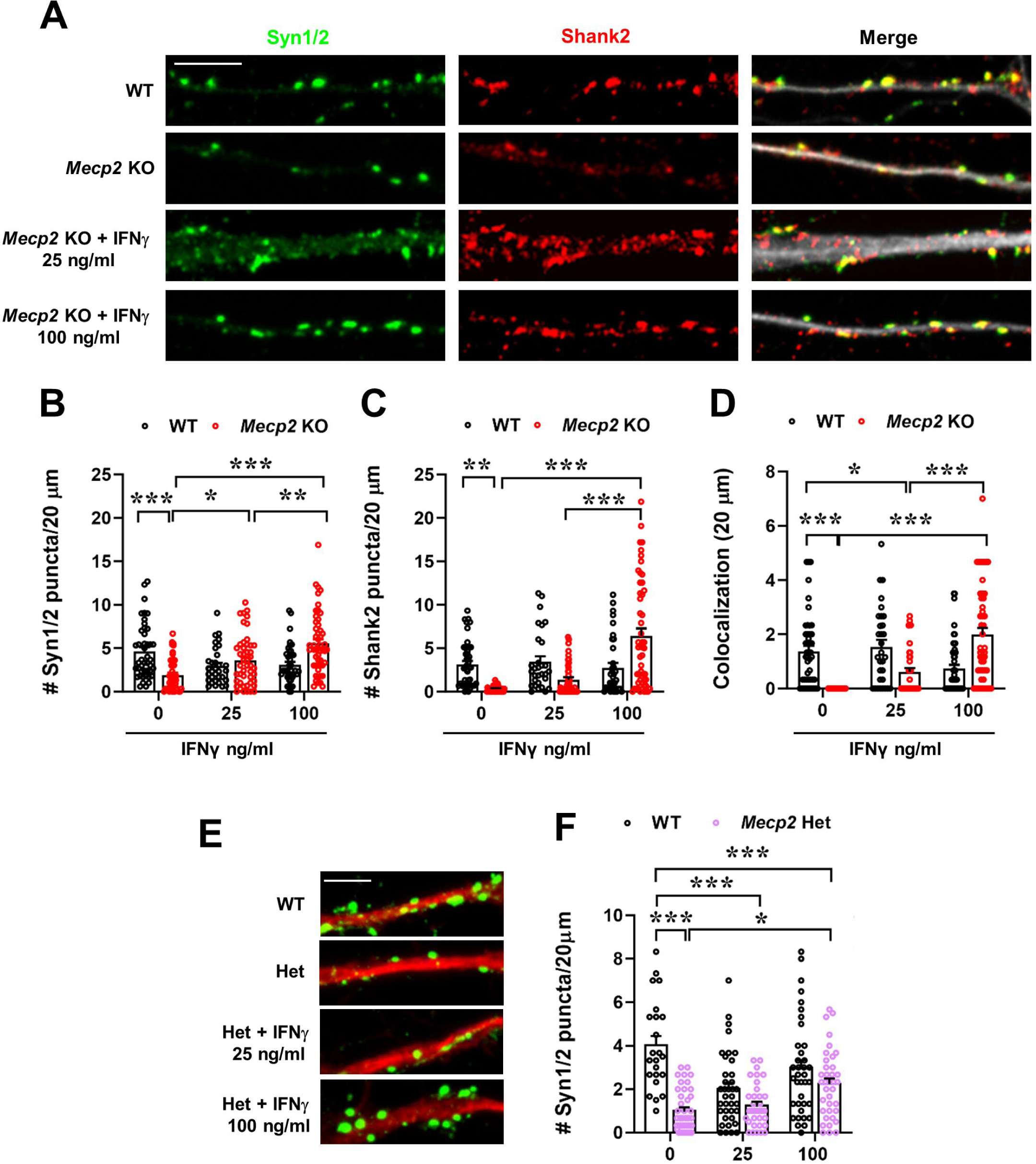
Synaptic defects of *Mecp2* KO cortical neurons are significantly improved by IFNγ treatment. **A)** Representative images of WT and *Mecp2* KO neurons (DIV14) untreated or treated for 24 hours at DIV13 with IFNγ (25 and 100 ng/ml) and immunostained for Synapsin1/2 (Syn1/2; green), Shank2 (red) and Map2 (white). **B-D)** Histograms indicate the mean ± SEM of the number of puncta counted in 20 µm for Synapsin1/2 (C), Shank2 (D) and colocalized puncta (E). *p<0.05, **p<0.01, ***p<0.001 by two-way ANOVA followed by Tukey post hoc test. **E)** Representative images of WT and *Mecp2* Het neurons (DIV14) untreated or treated for 24 hours at DIV13 with IFNγ (25 and 100 ng/ml) and immunostained for Synapsin1/2 (Syn1/2; green) and Map2 (red). **F)** The histogram indicates the mean ± SEM of Synapsin1/2 puncta density. *p<0.05, ***p<0.001 by two-way ANOVA followed by Tukey post hoc test. n=23 WT, n=37 WT+ IFNγ 25ng/ml, n=37 WT+ IFNγ 100ng/ml, n=49 *Mecp2* Het, n=35 *Mecp2* Het+ IFNγ 25ng/ml, n=37 *Mecp2* Het+ IFNγ 100ng/ml. Neurons derived from at least 3 different animals per genotype.

Eventually, we verified IFNγ efficacy *in vivo*. *Mecp2* KO symptomatic mice and WT littermates (P45) were i.c.m. injected with recombinant IFNγ (20 ng/ml) or an equal volume of vehicle. After 24 hours, a cohort of mice was tested for memory functions, a second one for motor abilities (**Fig 8A**). By NOR test, we reported that IFNγ significantly rescued memory deficits in KO mice (**Fig 8B**). Similarly, by Rotarod, we found that defects in motor learning and coordination were completely reverted by the acute injection of the cytokine (**Fig 8C,D**). On the contrary, analysis of respiratory alterations by whole-body plethysmography reported that an acute injection of IFNγ was unable to ameliorate defects in the frequency of respiration and inspiration/expiration time in KO mice (**Fig 8E-H**).

**Figure 8.**
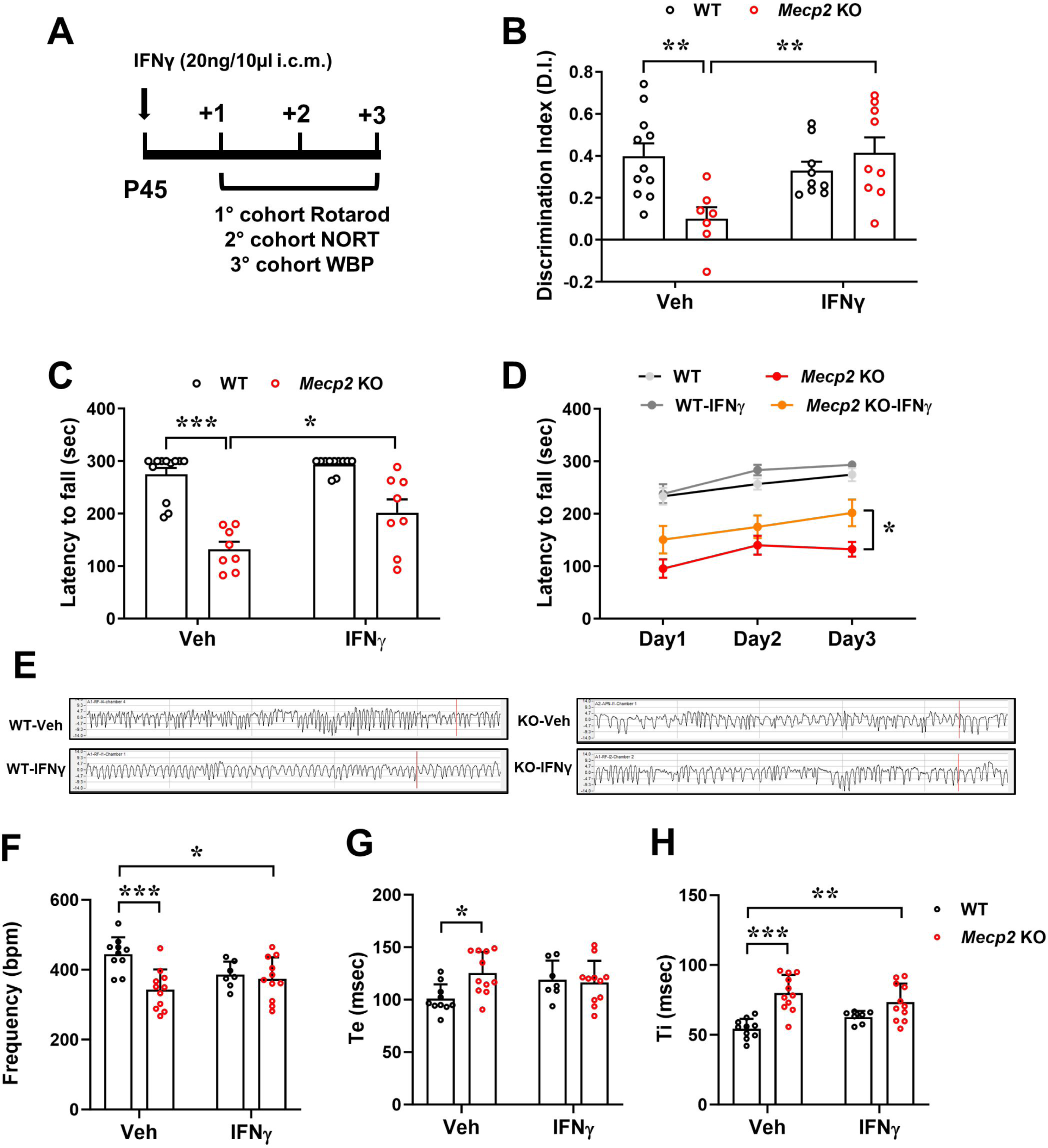
IFNγ injection in *Mecp2* KO mice reverts motor and cognitive impairments. **A)** Schematic representation of the *in vivo* experiments, in which IFNγ (20ng/10 µl) or vehicle (Veh) was stereotaxically injected in WT and *Mecp2* KO mice (P45; i.c.m.). Three cohorts of animals were used for assessing cognitive, motor and breathing defects. **B)** The histogram represents the mean ± SEM of the discrimination (D.I.) index values, assessed by novel object recognition (NOR) test. **p<0.01 by two-way ANOVA followed by Tukey post hoc test. n=11 WT-Veh, n=7 *Mecp2* KO-Veh, n=9 WT-IFNγ, n=9 *Mecp2* KO-IFNγ **C-D)** The graphs represent the mean ± SEM of the time (in seconds) spent on the rod during each day of the test, reflecting motor learning abilities (C). In D, the histogram shows the mean ± SEM of the time (in seconds) spent on the rod at the 3^rd^ day of the test. *p<0.05, ***p<0.001 by two-way ANOVA followed by Tukey post hoc test. n=10 WT-Veh, n=11 KO-Veh, n=7 WT-IFNγ, n=11 KO-IFNγ **E)** Representative traces from the WBP analysis of WT and KO mice, treated with vehicle or IFNγ. **F-H)** The histograms show the mean ± SEM of the frequency, time of expiration (Te) and time of inspiration (Ti) measured by whole-body plethysmography in freely-moving WT and KO mice, 72 hours after injection.

## Discussion

Thanks to the ability to promote damage repair and restore brain functions, stem cell therapy represents a promising therapeutic option for the treatment of many neurodegenerative diseases (Lindvall and Kokaia, 2010). Although several stem cell types are available for therapeutic treatments, neural progenitor cells (NPCs) offer the advantages of being highly stable in terms of self/renewal, expansion, and differentiation, and, importantly, do not originate tumors *in vivo* (Foroni et al., 2007). However, it was then proved that adult stem cells, including NPCs, can induce cancer when transplanted in a microenvironment different from their original one, thereby raising serious safety concerns (Melzi et al., 2010). These results reinforced the importance of treating neurological disorders with homotypic adult NPCs. Accordingly, several clinical trials based on NPC transplantation have been conducted so far (Genchi et al., 2023; Fan et al., 2023).

Although NPCs can differentiate into neurons, astrocytes and oligodendrocytes, their benefic effects upon transplantation are not exclusively based on cell replacement. Indeed, transplanted NPCs often maintain an undifferentiated phenotype and, by sensing the encountered microenvironment, engage into the release of different therapeutic molecules, including immunomodulatory, trophic and neuroprotective factors (Lindvall and Kokaia, 2010; Goldberg et al., 2015; Bacigaluppi et al., 2016; De Feo et al., 2017). Therefore, through a paracrine action exerted by a disease-adapted secretome, NPCs can influence many critical biological processes, such as survival, proliferation, differentiation and immune regulation (Drago et al., 2013, Willis et al., 2020).

While literature supporting the effectiveness of stem cells, and in particular of NPCs, for neurodegenerative disorders is wide (Lindvall and Kokaia, 2010), only few evidence are available regarding their application for neurodevelopmental diseases (NDDs) (Siniscalco et al., 2018; Donegan and Lodge, 2020). Most of these studies investigated the therapeutic efficacy of mesenchymal or hematopoietic stem cells on animal models of autism spectrum disorders (ASD) and in a limited number of cases amniotic epithelial or adipose- and urine-derived stem cells were used (Nabetani et al., 2023). Concerning RTT, the ability of mesenchymal stem cells to increase neurogenesis and the number of synapses in the brain of *Mecp2* KO mice was recently reported; however, stem cells were engineered to constantly produce BDNF, becoming a supplier system for the neurotrophin (Kim et al., 2021). Further, indications of a positive effect of NPC transplantation for NDDs can be deduced from a Chinese publication in which human NPCs (hNPCs) derived from aborted fetal tissues were administered to 13 children with RTT and 9 with ASD (Liu et al., 2013).

This evidence, together with the well-known widespread neurobiological abnormalities featured by the RTT brain, that could trigger NPCs to secrete neuroprotective molecules, prompted us to investigate the therapeutic effect of NPCs in RTT mouse models.

Importantly, *in vitro* we proved the existence of a beneficial crosstalk between NPCs and RTT neurons, that is not established with WT cells. This crosstalk induces NPCs to release factors able to rescue synaptic defects. These results led us to assess the value of NPC transplantation in the *Mecp2* null male brain that is characterized by the early onset of robust phenotypes that phenocopy several functional, cellular, and molecular features observed in human RTT samples (Guy et al., 2007; Cobolli Gigli et al., 2006). Coherently with previous pre-clinical studies performed in mouse models of neurodegenerative disorders, only few grafted cells were retrieved in the transplanted brain and a progressive decline of their number was observed. These cells did not enter in the parenchima but remained on the meninges proximal to the site of injection, maintaining an undifferentiated morphology and expressing the stem cell marker Nestin, thereby suggesting that any positive effect should depend on a bystander mechanism.

The therapeutic efficacy of NPCs for RTT emerged from the motor behavioral improvements and the partial amelioration of the cognitive functions of NPC-treated KO mice. Importantly, lifespan in KO animals was also significantly prolonged by treatment, although transplanted cells have a limited residency in the brain. We hypothesize that a longer and possibly a more robust benefit might result from repetitive cell transplantations. The potential of NPC transplantation was also tested in *Mecp2* Het female animals, which closely mimic the genetic condition of RTT patients. Rescued memory functions were displayed by Het mice upon transplantation, whereas no improvement was observed in motor abilities. We believe that several factors might have contributed to the observed limited efficacy, including i) the reduced grafting of NPCs; ii) the higher dispersion of results that typically derives from heterozygous mice; iii) the necessity of optimizing the time window for the treatment, since the progressive worsening of RTT symptomatology in heterozygous females is slower compared to KO animals (Ribeiro and MacDonald, 2020). To explore the mechanisms involved in the neuroprotective effects exerted by NPCs in RTT animals, we investigated the transcriptional changes induced by transplantation, highlighting the activation of immune system. Among the numerous deregulated pathways associated with immune response, we identified IFNγ as a strong candidate. Indeed, IFNγ is a soluble molecule, mainly secreted by immune cells, that can be easily targeted for therapeutic purposes (Monteiro et al., 2017). Further, this cytokine has been implicated in neuropsychiatric disorders (Schmidt et al., 2014; Arolt et al., 2000) and was already used in clinical settings as an immunomodulatory drug (Miller et al., 2009). Contrasting data are available regarding the effects of IFNγ on neurological symptoms, with some evidence reporting neurotoxic effects and other indicating neuroprotective ones (Mizuno et al., 2008; O’Donnel et al., 2015). This heterogeneity of data might depend on the source of the cytokine, the activated pathway and the targeted pathology. Of relevance, the capacity of an acute treatment with IFNγ to rescue social behavior defects in immuno-suppressed mice was recently proved (Filiano et al., 2016).

In RTT, reduced IFNγ concentration was demonstrated in patients’ serum (Leoncini et al., 2015), but single cell sequencing analysis of meningeal immune cells reported an increased IFNγ expression in KO mice (Li et al., 2023). While our data demonstrate that NPCs strongly activate IFNγ response pathway in the cerebellum of KO animals, they do not provide details regarding the source of the cytokine, since both NPCs and resident immune cells might concur to the activation of the cytokine pathway (Cossetti et al., 2014; Monteiro et al., 2017). Similarly, we do not currently know which cells respond to the cytokine. However, *in vitro* results reporting the activation of Stat1 upon NPC exposure and the effectiveness of IFNγ to ameliorate synaptic defects prompt to indicate neurons as target cells also *in vivo*.

In accordance with its ability to ameliorate synaptic alteration both in KO and Het neurons, a single injection of IFNγ in the CSF of symptomatic null animals completely reverted motor and memory impairments, which manly rely on alterations of synaptic plasticity. In contrast, respiratory alterations were not improved by the acute treatment, suggesting that an amelioration of respiratory defects might require the stabilization of less plastic molecular mechanisms.

In conclusion, we demonstrated that, by sensing the pathological milieu, NPCs secrete factors that rescue neuronal defects of RTT neurons and ameliorate neurological abnormalities of *Mecp2* deficient mice. Although the molecular mechanisms underpinning these beneficial effects might be different between *in vitro* and *in vivo* settings, transcriptomic analysis in the cerebellum of null mice after cell transplantation strongly indicated the possible involvement of IFNγ. Indeed, treatment with IFNγ of RTT neurons or *Mecp2* null animals ameliorated morphological and behavioral defects, respectively. Although we are aware that many other factors might mediate the beneficial effects of NPCs on RTT models, we believe that a preclinical study assessing the therapeutic efficacy of a prolonged treatment of IFNγ in RTT deserves further investigations.

## Methods and Materials

### Animals

The experiments were performed on the *Mecp2*^tm1.1Bird^ mouse model in the outbred CD1 genetic background, generated by crossing *Mecp2* heterozygous (Het) females in C57BL/6 background (B6.129P2(C)-Mecp2^tm1.1Bird^/J) with CD1 wild-type (WT) male mouse and maintaining animals on a clean CD1 background. CD1 *Mecp2* mutant line recapitulates the typical phenotype of *Mecp2* mutant animals in C57BL/6 background, with the advantage of higher breeding success and larger litters therefore facilitating basic and translational studies (Cobolli Gigli et al., 2016). For Neural Precursor Cells preparation, adult female C57BL/6 animals were used. Both CD1 WT male mice and C57BL/6 WT females were purchased from Charles River Laboratories. Mouse genotype was determined by PCR using the following primers: 5’-ACCTAGCCTGCCTGTACTTT-3’ forward primer for null allele; 5’ GACTGAAGTTACAGATGGTTGTG-3’ forward primer for wild type allele; 5’ CCACCCTCCAGTTTGGTTTA-3’ as common reverse primer.

For neuronal cultures, WT and *Mecp2* mutant embryos were generated by mating Het females with WT mice. The day of vaginal plug was considered E0.5 and primary neurons were prepared from E15.5 embryos. For *in vivo* and *ex vivo* experiments, *Mecp2* KO animals (P45/47) and *Mecp2* Het mice (P180), and the corresponding WT littermates, were randomly assigned to the treatment groups. Animals were sacrificed by rapid decapitation or by transcardiac perfusion depending on experimental needs.

Animals were housed in a temperature- and humidity-controlled environment in a 12 h light/12 h dark cycle with food and water *ad libitum*. All procedures were performed in accordance with the European Union Communities Council Directive (2010/63/EU) and Italian laws (D.L.26/2014). Protocols were approved by the Italian Council on Animal Care in accordance with the Italian law (Italian Government decree No. 175/2015-PR, No. 210/2017 and No. 187/2022-PR).

### Cell cultures

#### Neural Precursor Cells (NPCs)

Adult female C57BL/6 mice (6 to 8 weeks old, 18-20 gr) were anaesthetized by an intraperitoneal injection of Tribromoethanol (250 mg/Kg, i.p.) and the brain was removed and positioned in sterile PBS. Brain coronal sections were taken 3 mm from the anterior pole of the brain, excluding the optic tracts and 3 mm posterior to the previous cut. The subventricular zone (SVZ) of the lateral ventricles was isolated from the coronal section using iridectomy scissors. Tissues derived from at least two mice were pooled and digested for 30 minutes at 37°C with Earl’s Balanced Salt Solution (EBSS) (#E2888, Sigma-Aldrich) containing 200 mg/l L-Cysteine (#C7352, Sigma-Aldrich), 200 mg/l EDTA (#E6511, Sigma-Aldrich), 2U/ml Papain (#P4762, Sigma-Aldrich) (Pluchino et al., 2003). Sample was centrifuged at 200 g for 12 minutes, the supernatant was removed a nd the pellet was mechanically disaggregated with 2 ml of EBSS. The pellet was centrifuged again at 200 g for 12 minutes and then dissociated with a pipette. Cells were plated in Neurocult proliferation medium, containing Neurocult Basal Medium (#05702, Stem Cell Technologies), Proliferation Supplement (#05701, Stem Cell Technologies), 10 ng/ml FGF (#1370 9505 00, Provitro), 20 ng/ml EGF (#1325 0510 00, Provitro), 0.0002% Heparyn (#H3393-100KU, Sigma-Aldrich) and 1% Penicillin/Streptomycin (P/S; #P0781, Sigma-Aldrich). After approximately one week, a small percentage of the isolated cells begun to proliferate, giving rise to neurospheres, which grow in suspension. Neurospheres were centrifuged at 50 g for 10 minutes, the supernatant removed, 200 ml of Accumax (#A7089, Sigma-Aldrich) was added and the tube was incubated at 37°C for 10 minutes. The pellet was mechanically dissociated and cells were plated at a density 7500 cells/cm^2^ for passages or used for *in vitro* and *in vivo* experiments.

For *in vivo* experiments, NPCs were infected with 3 x 10^6^ T.U./ml of a third-generation lentiviral vectors. Cells were dissociated 12 hours before infection and plated at high density (1.5 x 10^6^ cells in a 75 cm^2^ flask) in 10 ml of medium. 48 hours after infection, cells were harvested, centrifuged at 200 g for 12 minutes and plated without dissociation at a 1:1 ratio. After 3 passages *in vitro*, FACS analysis was performed to verify the efficiency of the infection (Pluchino et al., 2005). All cells used were tested and were negative for mycoplasma.

#### Primary cortical neurons

At E15.5, WT, KO and Het mouse embryos were used to prepare neuronal cultures (Frasca et al., 2020). Brains were removed under a microscope and immersed in ice-cold Hank’s Buffered Salt Solution (HBSS; #14175-095, Life Technologies). Meninges were gently removed, cerebral cortex was rapidly dissected and maintained in ice-cold HBSS. Tissues were incubated with 0.25% trypsin/EDTA (# 25200-056, Life Technologies) for 7 min at 37°C and the digestion was blocked with 10% FBS (#10500064, Gibco) in DMEM (#41966-029, Life Technologies). Cortices were then mechanically dissociated by pipetting in DMEM, containing 10% FBS, 1% L-glutamine (#G7513, Sigma-Aldrich), 1% P/S. Neurons were seeded in neuronal medium [Neurobasal (#21103049, Gibco), 2% B27 (#A3582801, Gibco), 1% L-Glutamine, 1% P/S] on coated glass coverslip (Neuvitro) in 24-well plates (40000 cells/well) for immunofluorescence analysis, in poly-D-lysine (0.1 mg/ml; #P7886, Sigma-Aldrich)-coated 6-well plates (200,000 cells/well) for western blot experiments and in 96-well plates (10,000 cells/well) for MTT assay (#M2003, Sigma-Aldrich).

To facilitate neuronal morphological analysis, by in-utero electroporation we transfected some cortical neurons in embryos with a GFP expressing plasmid. Timed-gestation Het females (E13.5) were deeply anesthetized with Tribromoethanol (250 mg/Kg; i.p.) and uterine horns were exposed by midline laparotomy. Plasmid DNA (pCAG vector with an iresGFP empty; 0.5 μg/μl of DNA) was injected in the telencephalic ventricle using a pulled micropipette. Then platinum electrodes were placed outside the uterus over the telencephalon and 4 squared pulses of 40 V were applied at 50 ms intervals. The uterus was then placed back in the abdomen; muscle and skin were closed with sutures.

#### NIH3T3 fibroblast cultures

NIH3T3 fibroblasts were maintained in growth medium (DMEM supplemented with 10% FBS, 1% L-glutamine, 1% P/S) in T75 flasks at 37°C with 5% CO_2_. When confluent, cells were washed with PBS (#ECB4004L, Euroclone) and then incubated with trypsin (#ECB3052D, Euroclone) for 3-5 minutes at 37°C. Trypsin was inactivated with growth medium and cells were then centrifuged at 1200 g for 7 minutes. The supernatant was removed, and the pellet was mechanically disaggregated with 1 ml of medium.

#### NPCs/NIH3T3-neuron co-cultures

NPCs and NIH3T3 fibroblasts were seeded in neuronal medium on transwell inserts (pore size = 0,4 µm; Corning Costar), at a density 1:100 respect to neurons. When neurons attached to coverslips or wells (∼ 2 hours after plating), inserts with NPCs and NIH3T3 cells were carefully transferred on neurons and maintained until DIV7 or DIV14, without changing the medium.

From co-cultures between neurons and NPCs/NIH3T3, conditioned medium (CM) was collected at DIV14. CM was immediately centrifuged at 1200 rpm for 5 minutes at 4 °C to remove cellular debris and stored at - 80°C until use.

#### Neuronal treatment

For the treatment of neurons with CM, the day of the experiment, CM samples were left at room temperature (RT) for 10 minutes, then heated at 37°C in a water bath for 5 minutes and finally added to neurons, in a ratio 1:1 respect to neuronal culture medium. CM treatment was conducted for 24 hours (DIV13-DIV14). To treat neurons with IFNγ, WT and *Mecp*2 deficient neurons were treated with 25, 75 or 100 ng/ml of mouse recombinant IFNγ (GRF-15448; Immunological Sciences) or nuclease-free-H_2_0 (vehicle) as control for 24 hours (DIV13-DIV14). IFNγ was dissolved in sterile H_2_0, sub-divided in 5µl-aliquots and stored at −20°C.

### Intra-cisterna magna injection

Mice were anesthetized with a mixture of O_2_ and 3% isofluorane and during surgery anaesthesia was maintained at 1.5% isofluorane and respiration continuously monitored. Mice were fixed on a stereotactic device (David Kopf Instruments). Vehicle (10 µl of sterile PBS) or NPCs (10^6^ cells/10 µl) were injected intra cisterna magna (i.c.m.) in WT, *Mecp2* KO and Het mice. IFNγ administration was performed in WT and KO animals. A 10 µl Hamilton syringe was placed between the atlas and occipital bone at 35°, and advanced to puncture the cisterna magna. The following stereotactic coordinates were used: *x* = 0; *y* = –7.5 mm from λ; *z* = –4.5 mm from muscle plane. A total volume of 10 μl was injected in a 5-min time period and the needle was placed in situ for 3 minutes after injection before being slowly removed. Since NPCs were injected in CD1 animals (allogenic transplant), mice were treated with subcutaneous Ciclosporin A (15 mg/Kg, s.c; Novartis), starting the day before transplantation and for the consecutive 15 days. PBS-injected WT/KO mice were similarly treated with the immunosuppressive drug.

### Behavioural Analyses

#### Phenotypic characterization

*Mecp2* KO, Het mice and WT littermates were tested for the presence or absence of RTT-like symptoms with a previously described scoring system (Guy et al. 2007; Cobolli-Gigli et al. 2006; Gandaglia et al., 2018; Scaramuzza et al., 2021). At each session, an observer blind to the treatment assigned a score for general condition, mobility, gait, hindlimb clasping and tremor. Importantly, when the sum of the individual scores was >8, the mouse was euthanized for ethical reasons. Day of euthanasia was considered day of the death, without distinguishing it from natural death. Graphically, for each parameter the evolution of symptomatology was represented by a cumulative plot, obtained by plotting for each day the score summed to all preceding ones.

#### Rotarod Test

Rotarod test was set in a three-day paradigm using a 5 lane Ugo Basile’s Rotarod for mice. Every day, each mouse was subjected to three trials. For each trial, animals were placed on the rotating rod (4 rpm) and were allowed to habituate for 10 seconds. Then the rotation speed was gradually accelerated every 30 seconds across a period of 300 seconds, from 4 rpm to 40 rpm. For each mouse, trial terminated when animal fell of the rod or achieved the maximum time (300 seconds) (Buitrago et al., 2004). Rotarod test was performed 10 days after NPCs/vehicle injection, and 24 hours after IFNγ/vehicle injection.

#### Novel Object Recognition test

The novel object recognition (NOR) test was performed in a square arena of 45 x 45 cm (Balducci et al., 2017; Scaramuzza et al., 2021). On day 1, mice were first allowed to habituate to the testing arena in a 10 minutes session and mobility in the open field was monitored. On day 2, animals underwent the training phase (5 minutes), in which two identical objects were introduced into the arena, allowing the mouse to explore them. Finally, after 4 hours from the training phase, mice were tested for their memory (5 minutes). The discrimination index (D.I.), defined as the difference between the exploration time for the novel object and the familiar object, divided by total exploration time, was calculated. The sessions were recorded with the video tracking software EthoVision XT 14 (Noldus).

#### Plethismography

The breathing activity of freely-moving mice was recorded in a Vivoflow whole-body plethysmography system (EMKA Technologies) using a constant flow pump connected to the animal chamber, thus ensuring proper inflow of fresh air. Four plethysmography chambers of 200 ml, calibrated by injecting 1 ml of air, were used for simultaneous measurements. Breathing cycles were recorded for 15 minutes under normal-ventilation after an adaptation phase of 10 min in the recording chamber during the previous 5 days. The signal was amplified and recorded with IOX2 software.

### Immunofluorescence

For immunofluorescence analysis on brain sections, animals were anesthetized with Tribromoethanol (250 mg/ml, i.p.) and transcardially perfused with 30 ml ice-cold PBS, followed by 50 ml 4% paraformaldehyde (PFA). Brains were removed and post-fixed in 4% PFA for additional 24 hours at 4°C, then cryoprotected for 48 hours at 4°C in 30% sucrose in PBS and frozen in n-pentane at −30°C for 3 minutes. Coronal sections were cut with a cryostat (Leica Biosystems) along the rostro-caudal orientation of the brain and mounted on SuperFrost Plus microscope slides (Menzel-Glaser) or, alternatively, put on a multiwell in PBS supplemented with NaN_3_.

Immunofluorescence on serial brain sections mounted on slides (20 μm in thickness) was performed to study GFP-positive cells distribution; free-floating immunofluorescence was conducted on slices (30 μm in thickness) to study GFP-positive cells differentiation. Brain sections were washed three times for 5 minutes in PBS and incubated for 1 hour in blocking solution [10% normal goat serum (NGS; #50197Z, Invitrogen), 0.1% Triton X-100 in PBS]. Sections were then incubated overnight at 4°C with the primary antibody diluted in 1% NGS, 0.1% Triton X-100 in PBS. The following primary antibodies were used: anti-GFP (1:500; #A10262, Invitrogen), anti-GFAP (clone GA5, 1:1000; #MAB 3402, Millipore), anti-Olig2 (1:200; #ab136253, Abcam), anti-NeuN (clone 60, 1:100; #MAB377, Millipore) and anti-Nestin (clone rat-401, 1:100; #MAB353, Immunological Science). Slices were washed 3 times for 10 minutes in PBS and then incubated for 1 hour with the Alexa-Fluor secondary antibody diluted 1:500 in blocking solution. Sections were washed 8-10 times in PBS for 5 minutes and incubated with DAPI (0.1 mg/ml in PBS; #62248, Invitrogen) to stain nuclei; sections were washed in PBS and finally mounted with the Fluoromount mounting medium (#F4680; Sigma-Aldrich). Images were acquired at Nikon Ti2 Microscope equipped with an A1+ laser scanning confocal system and a 100x oil-immersion objective. To estimate the number of transplanted NPCs in KO brains, GFP^+^ cells were manually counted on serial brain sections under an epifluorescence microscope (Nikon Eclipse Ti). The number of cells per section and plane was multiplied by the number of sections in between the sections, thus obtaining overall numbers of GFP^+^ NPCs per brain (Bacigaluppi et al., 2016).

For immunofluorescence on cultured cells, cortical neurons (DIV7 or DIV14) seeded on glass coverslips were fixed for 8 minutes with 4% PFA dissolved in PBS with 10% sucrose, then washed three times with PBS and stored in 0.1% NaN_3_ in PBS at 4°C. Cells were permeabilized in 0.2% Triton X-100 in PBS for 3 minutes on ice. Then, cells were washed in 0.2% BSA in PBS, then in blocking solution (4% BSA in PBS) for 15 minutes and finally incubated with primary antibodies overnight at 4°C. Primary antibodies were diluted in 0.2% BSA in PBS as follow: anti-Map2 (clone D5G1, 1:1000; #8707, Cell Signalling), anti-Synapsin1/2 (1:500; #106006, Synaptic System), anti-Shank2 (1:300; #162211, Synaptic System). After washing in BSA 0.2% in PBS, cells were incubated with the specific Alexa Fluor secondary antibody (1:500; Thermo Fisher) in 0.2% BSA in PBS for 1 hour at RT. After 5 washes in PBS, nuclei were stained with DAPI and cells were washed in PBS. Lastly, glass coverslips were mounted on microscope slides with Fluoromount mounting medium and stored at 4°C until image acquisition.

### Analysis of neuronal morphology and synaptic markers

GFP^+^ neurons were acquired at DIV7 at epi-fluorescence microscopes (Nikon Eclipse Ti) using an excitation wavelength of 488 nm with a 20x objective. To characterize neuronal morphology, images were processed using ImageJ software. Sholl analysis plugin was used to study the complexity of the dendritic arbour and NeuronJ plugin to measure dendritic length (Patnaik et al., 2020; Baj et al., 2014). Since both plugins require binary masks, we manually traced all the GFP^+^ dendrites with Photoshop for each neuron and obtained images were binarized. We analyzed dendritic arborization by performing Sholl analysis using a radius step size of 10 µm. The same reconstructed neurons were used to analyze the total lengths of dendrites by NeuronJ.

To analyze synaptic puncta density and colocalization, Z-stacks images (127.28 × 127.28 µm^2^, 1,024 × 1,024-pixel resolution, 16-bit greyscale depth) were acquired at Nikon Ti2 Microscope equipped with an A1+ laser scanning confocal system and a SR Apo TIRF 100x oil-immersion objective, using a step size of 0.3 µm. For each dataset, images were acquired in four channels (laser wavelength for DAPI: 409.1 nm; laser wavelength for Synapsin1/2: 487.5 nm; laser wavelength for Shank2: 560.5 nm; laser wavelength for Map2: 635.5 nm) and parameters were maintained constant within the experiment (offset background, digital gain, laser intensity, pinhole size, scanning speed, digital zoom, scan direction, line average mode).

Puncta density was calculated by counting synaptic puncta within a manually selected ROI (length 20 µm on 3 primary branches/neuron) by ImageJ software. Only puncta with a minimum size of 0.16 µm^2^ were counted using *Analyze Particles*. To assess puncta co-localization of pre- and post-synaptic markers, the plugin *Colocalization highlighter* was run on each Z-stack image. Colocalized puncta were quantified in manually selected ROIs of the binary mask created from the maximum intensity projection. Only puncta with a minimum size of 0.1 µm^2^ were counted (Frasca et al., 2020; Albizzati et al., 2023).

### Electrophysiological measurements

Whole-cell patch-clamp recordings were obtained from WT and *Mecp2* Het cortical neurons at DIV14 in the voltage-clamp modality using the Axopatch 200B amplifier and the pClamp-10 software (Axon Instruments). Recordings were performed in Krebs’-Ringer’s-HEPES (KRH) external solution (NaCl 125 mM, KCl 5 mM, MgSO_4_ 1.2 mM, KH_2_PO_4_ 1.2 mM, CaCl_2_ 2 mM, glucose 6 mM, HEPES-NaOH pH 7.4 25 mM). Recording pipettes were fabricated from glass capillary (World Precision Instrument) using a two-stage puller (Narishige); they were filled with the intracellular solution potassium-gluconate (KGluc 130 mM, KCl 10 mM, EGTA 1 mM, HEPES 10 mM, MgCl_2_ 2 mM, MgATP 4 mM, GTP 0.3 mM) and the tip resistance was 3-5 MΩ. In order to identify excitatory postsynaptic currents in miniature (mEPSCs), cortical neurons were held at −70 mV and Tetrodotoxin (TTX) 1 μM was added to the external solution. The recorded traces have been analyzed using Clapfit-pClamp 10 software, after choosing an appropriate threshold.

### Western Blot

Neurons at DIV14 were washed briefly with sterile PBS and collected in 15% 2-mercaptoethanol in sample buffer. Samples were heated at 95°C for 5 minutes and then separated on a SDS-PAGE on TGX-stain free precast gel (4-15% of acrylamide gradient) and transferred on a nitrocellulose filter using a semi-dry transfer apparatus (TransBlot SD; Bio-Rad). Then, membranes were incubated for 1 hour in blocking solution [5% BSA in 0.1% Tween-20 in Tris-buffered saline containing (TBST)], and incubated overnight at 4°C with the following primary antibodies: rabbit anti-phosphorylated Stat1 (clone 58D6, 1:1000; #9167, Cell Signalling) or rabbit anti-Bdnf (1:1000; ab108319, Abcam). After three washes in TBST, membranes were incubated with HRP-conjugated secondary antibody for 1 hour at RT (1:10000; Jackson ImmunoResearch). The immunocomplexes were visualized by using the ECL substrate (Cyanagen) and Essential V6 imaging platform, UVITEC system (Cleaver Scientific Ltd). Band density measurements were performed using UVITEC software. Results were normalized to total protein content visualized by a TGX stain-free method (Bio-Rad).

### RNA extraction, qRT-PCR, RNA Sequencing and bioinformatics

Twenty days after transplantation, WT and *Mecp2* KO animals were euthanised by cervical dislocation and brain quickly removed. Total RNA from cerebella was extracted using Purezol (Bio-Rad) and quantified using a NanoDrop spectrophotometer. RNA integrity was assessed by *RNA 6000 Nano Reagent kit* on a Agilent 2100 Bioanalyzer (Agilent Technologies) (Carli et al., 2021; Albizzati et al., 2022). All samples showed an RNA integrity number (RIN) >7.5.

RNA-sequencing was conducted by the Genomic Facility of the IRCCS San Raffaele Hospital (CTGB). cDNA library of the collected RNA samples was obtained using the TruSeq RNA Library Prep kit from Illumina. Single-end RNA Sequencing was performed on an Illumina HiSeq 2500 Next Generation Sequencing instrument. The quality of the reads was verified with FastQC (v.0.11.8) (Andrews, 2010). Sequencing adapters were trimmed with Trimmomatic (v. 0.39) (Bolger et al., 2014) and the trimmed fastq files were aligned to the reference genome with STAR (v. 2.53a) (Dobin et al. 2013). The mouse reference genome used for the alignment was the Mus musculus GENCODE release M22 (GRCm38.p6). Finally, the mapped reads were counted, grouped by genes, with featureCounts (v. 1.6.4) (Liao et al., 2014) setting the strandness of the single-end reads as ‘reverse’. Quality check of the different steps of the analysis was performed with MultiQC (Ewels et al., 2016) (**Fig EV4**). Data are deposited in ArrayExpress (E-MTAB-12813). Principal Component Analysis was performed using the prcomp function in R, using the 500 most variable genes in term of Reads Per Kilobase Million (RPKM).

Differential gene expression analysis was performed with the DESeq2 bioconductor package (Love et al., 2014) on the following comparisons: *Mecp2* KO *versus* WT, both treated with PBS as control (*Mecp2* KO-PBS, WT-PBS), to assess the effects of *Mecp2* deficiency; *Mecp2* KO treated with NPCs (*Mecp2* KO-NPCs) vs *Mecp2* KO-PBS, to assess the effects of the treatment on KO animals; WT treated with NPCs (WT-NPCs) vs WT-PBS, to assess the effects of the treatment on WT animals; *Mecp2* KO-NPCs vs WT-PBS, to assess the rescue on KO animals. A p-value adjusted with FDR <0.05 was used to determine the significance of DEGs.

Over Representation Analysis (ORA) (Boyle et al., 2004) was performed using the Bioconductor package clusterProfiler (Yu et al., 2012). The function ‘simplify’ was used to remove redundancy of enriched GO terms.

Gene Set Enrichment Analysis (GSEA) (Subramanian et al., 2005) (version 4.1.0) was performed using the R package fgsea (Korotkevich et al., 2021) on shrunken, log-normalized exonic fold changes from DESeq2.

Background was set to all expressed genes in this study and 1,000 permutations were set to generate a null distribution for enrichment score in the hallmark gene sets and functional annotation gene sets.

For qRT-PCR experiments, RNA was reversely transcribed using the RT^2^ First Strand Kit (#330404, Qiagen) as instructed by the manufacturer. The resulting cDNA was used as a template with SYBR Green Master Mix (Applied Biosystems) with designated primers (**Table 1**). Melting curve showed a single product peak, indicating good product specificity. Actin and Rpl13 were used as housekeeping genes and geometric average was calculated and used for the analysis of fold change in gene expression using the 2(-delta Ct) method.

**Table 1.**
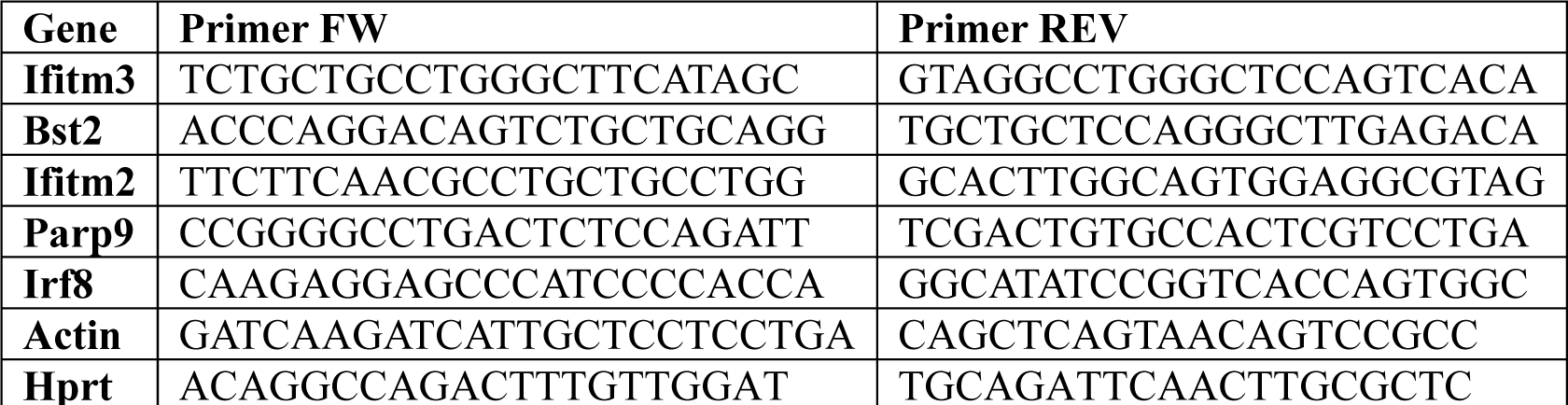
List of primers used for qRT-PCR.

### Statistical analysis

All data are expressed as mean ± SEM. Before any statistical analysis, normality distribution was assessed for each dataset by D’Agostino and Pearson tests. Outliers were evaluated by ROUT test (Q=1%) or Grubb’s test (α=0.05%). Statistical significance for multiple group comparisons was determined by one-or two-way analysis of variance (ANOVA), followed by Tukey post hoc test. Mann–Whitney tests was used for comparing two groups. All statistical analyses were performed using GraphPad Prism 9.

## Acknowledgements

This work was supported by the Italian parents’ association “PRO RETT Ricerca” to A.F and N.L., by Banca d’Italia, and Jerome Lejeune Foundation. We are very grateful to professor Elena Borroni from University of Milan for her generous gift of antibody against phosphorylated Stat1. We are grateful to all members of N.L. and G.M. laboratories for helpful discussions.

## Author contribution

A.F., F.M., E.B., M.I., M.B.A., F.M.P., A.P., F.B., L.P., C.C., F.A. performed the experiments and analysed the data. A.F and F.M. prepared the figures. A.F. and N.L. designed the study and wrote the manuscript. U.B. analysed transcriptomic data, performed bioinformatics analyses. G.M. assisted in interpreting and discussing the results. All the authors revised the manuscript.

## Conflict of interest

The authors declare that they have not conflict of interest.

## The Paper Explained Problem

*MECP2* mutations cause RTT, the first genetic cause of severe intellectual disability in girls worldwide. Although potentially treatable, so far only a drug mainly affecting fine motor skills and communication is available. The identification of valid therapy is hampered by the still limited comprehension of the neurological roles of MeCP2 and the consequences of its loss of function. NPC transplantation has already been proved safe and efficacious in many neurological disorders but a comprehensive study of NPC efficacy in RTT is lacking.

## Results

We have collected several data indicating that NPC transplantation improves RTT-like symptoms in *Mecp2* mutant animals and that NPC-secreted factors are responsible for the beneficial effects on *Mecp2* mutant neurons. Importantly, our study disclosed that the activation of the Interferon-γ pathway participates to the observed benefic effects; accordingly, we proved the therapeutic efficacy of this cytokine in RTT models.

### Impact

Willing to respond to the unmet need of identifying novel therapies for RTT, we proved the therapeutic potential of NPCs and we identified Interferon-γ as a possible novel healing molecule for RTT. Although this study remains at the preclinical realm, the obtained positive results represent the “proof-of-principle” required to directly inform new clinical trials design.

### For More Information

ProRett Ricerca https://prorett.org/ https://www.ebi.ac.uk/biostudies/arrayexpress

## Expanded View Figure legends

**Figure EV1. A)** Sholl analysis reports the capacity of NPCs to increase dendritic complexity also in WT neurons. The graph depicts the mean ± SEM of the number of intersections of WT neurons cultured alone, or with NIH3T3 or NPCs from DIV0 to DIV7. **B)** The histogram shows the mean ± SEM of the total number of intersections calculated by Sholl analysis for WT and KO neurons cultured alone, or in culture with NIH3T3 or NPCs from DIV0 to DIV7. *p<0.05, **p<0.01 by two-way ANOVA followed by Tukey post hoc test. **C)** Representative traces of excitatory postsynaptic current in miniature (mEPSCs) recorded in primary wt and Het neurons (left) and in Het neurons treated with NPC or NIH3T3 (right). **D**) Histograms represent the mean ± SEM of the mEPSCs frequency (Hz) and amplitude (pA) both expressed as values normalized on the frequency of WT neurons or amplitude of WT neurons. n= 26 WT; n= 24 Het; n= 19 Het + NPCs; n= 14 Het + NIH3T3. Data derived from 3 independent experiments. * p<0.05 by one-way ANOVA followed by Kruskal-Wallis test.

**Figure EV2. Phenotypic characterization of NPC-transplanted *Mecp2* KO mice. A-E)** Behavioral scoring was performed by a researcher blind to the treatment, assigning a score between 0 and 2 to general aspect (A), mobility (B), gait (C), tremor (D) and clasping (E). For each parameter, the graph reports its progression by a cumulative plot, in which the mean ± SEM of each value is obtained by summing the score of each day with those assigned the preceding days. Asterisks indicate a significant difference between KO-PBS and KO-NPCs; hashtags denote a difference between KO-NPC and WT mice. * or # p<0.05, ***p<0.001 by two-way ANOVA followed by Tukey post hoc test. n=15 WT-PBS, n=15 WT-NPCs, n=9 *Mecp2* KO-PBS, n=12 *Mecp2* KO-NPCs. **F)** Representative traces of the distance travelled during the first day of NOR test by WT-PBS, *Mecp2* KO-PBS, WT-NPCs and *Mecp2* KO-NPCs animals. The corresponding data are reported in Figure 3H.

**Figure EV3. Gene Ontology (GO) analysis. A, B)** Dot plot of Gene Ontology (GO) enriched pathway analysis in the cerebellum, indicating the top 30 most enriched pathways of the comparison between KO *versus* WT (A), and KO+NPC *versus* WT (B) samples.

**Figure EV4. Quality check of the RNA-seq experiment. A)** Table describing the genotype and the transplantation conditions of the sequenced samples. **B)** Dropped vs surviving reads after Trimmomatic adapter trimming. **C)** Unique reads number s duplicated reads. **D**) STAR alignment scores: number of uniquely mapped reads, mapped to many loci or unmapped. E) HTSeq count assignments to genes.

**Figure EV5. IFNγ does not affect neuronal survival and activates its downstream kinase *in vitro*. A)** The histogram represents the cell survival (% untreated WT neurons), assessed by MTT assay. IFNγ was added for 24 hours in DIV13 primary neurons and three doses were tested: 25, 75 and 100 ng/ml. **B)** The histograms reports the mean ± SEM of the levels of phosphorylated STAT1 after IFNγ treatment. Data are normalized to total protein content, visualized by a TGX stain-free technology. Representative bands of phosphorylated STAT1, and the corresponding lanes of TGX-stain-free gel, in WT and *Mecp2* KO neurons treated or not with IFNγ are depicted. § p<0.001 denotes a significant difference respect to the corresponding untreated control of the same genotype.

## Notes

### Competing Interest Statement

The authors have declared no competing interest.

